# Recurrent RNA-lipoplex vaccination is required to sustain functional tumor-infiltrating neoantigen-specific CD8 T cells and therapeutic efficacy

**DOI:** 10.1101/2025.11.16.687774

**Authors:** Justin T. Gibson, Milena Hornburg, Sophie Lehar, Vincent Javinal, Lisa Liao, Mark J. McCarron, Tamaki Nozawa Jones, Yoko Oei, Debra Dunlap, Kevin A. Marroquin, Amy A. Lo, Jan H. Bergmann, Ann-Jay Tong, Ariane M. Nissenbaum, Emily C. Freund, Aude-Hélène Capietto, Cecile C. de la Cruz, Ugur Sahin, Ira Mellman, Jill M. Schartner, Lélia Delamarre

## Abstract

Cancer vaccines induce durable, polyepitopic T cell responses, and show promising clinical benefit in adjuvant settings, yet they are largely ineffective in advanced disease. Using a clinically relevant RNA–lipoplex vaccine, we investigated the efficacy constraints in a preclinical model. Vaccination remodeled the tumor microenvironment (TME), increasing T cell infiltration and promoting a proinflammatory myeloid compartment. This was associated with complete regression of smaller, immature tumors, but only delayed growth of larger, established tumors. While vaccine-induced T cells were long-lived and functional in peripheral tissues, intratumoral T cells declined rapidly in abundance, diversity, and function, reverting to a prevaccine-like state. scRNA-seq suggested that this was driven by a pro-apoptotic program, with surviving T cells showing signatures of cellular stress and impaired activation. Importantly, recurrent vaccination replenished functional T cells in the TME and enhanced efficacy. These findings highlight the importance of optimizing vaccine schedules and tailoring therapeutic strategies to tumor stage.

## Introduction

Therapeutic vaccines targeting mutated cancer antigens, also known as neoantigens, are emerging as a promising treatment for patients with resected cancers. In a randomized phase 2 clinical trial, the mRNA-lipid nanoparticle (RNA-LNP) neoantigen vaccine mRNA-4157/v940 combined with the immune checkpoint inhibitor pembrolizumab prolonged recurrence-free survival in patients with high-risk resected melanoma in comparison with pembrolizumab monotherapy^1^. Similarly, results from a phase 1 study suggested that combination of the mRNA-lipoplex (RNA-LPX) neoantigen vaccine, autogene cevumeran, with atezolizumab checkpoint inhibitor and mFOLFIRINOX standard-of-care chemotherapy, induced neoantigen-specific CD8 T cells that may associate with delayed recurrence in patients with surgically resected pancreatic ductal adenocarcinoma (PDAC)^2,3^. While these results in the adjuvant setting following surgical resection are promising, the efficacy of neoantigen cancer vaccines has been more limited in patients with advanced solid tumors, despite the induction of substantial T cell responses^4–7^.

A number of factors may underlie the lack of clinical activity in patients with established disease. Following successful priming and expansion in secondary lymphoid organs, vaccine-induced T cells must traffic to and infiltrate into the tumor bed where they can carry out effector functions through recognition and killing of tumor cells^8^. It stands to reason that the magnitude of the vaccine-induced T cell response may be insufficient to eliminate bulky tumors. Infusion of billions of ex vivo-expanded autologous tumor-infiltrating lymphocytes (TILs) is needed to provide significant clinical benefit to patients with metastatic cancers^9–12^. Persistence and expansion of effector T cells in the tumor microenvironment (TME) is also critical for anti-tumor activity. This process can be limited by both chronic T cell stimulation and the hostile TME, leading to T cell exhaustion or activation-induced cell death (AICD). Recent findings also indicate that T cells require additional positive signals obtained from cross-talk with dendritic cells (DCs) to maintain their function and expand within tumors^13–16^ .

Tumors can evade the immune system by employing a wide array of mechanisms that limit immune cell infiltration, function, and recognition. Remodeling of the tumor vasculature and development of physical stromal barriers can function to limit T cell infiltration^17^. The TME, consisting of tumor cells, stroma-associated cancer-associated fibroblasts, and immune cells, including myeloid cells and regulatory T cells, suppresses T cell function through upregulation of surface inhibitory molecules (e.g. PD-L1) and production of immunosuppressive cytokines and metabolites (e.g. TGFb, IL-10, adenosine, and prostaglandin E2)^18–20^. Finally, tumor cells can escape T cell recognition by downregulating expression of major histocompatibility complex class I (MHCI) and expression and presentation of the target antigen^9,21–24^. In addition to active suppression of the immune response, rapid proliferation of tumor cells profoundly alters immune cell function through depletion of glucose, amino acids and oxygen. This nutrient depleted environment constrains metabolic pathways including glycolysis, oxidative phosphorylation, and lipid metabolism that are critical for T cell cytokine production, cytotoxic function and survival^25^.

To design superior vaccine platforms and more potent combination therapies for improving patient outcomes, it is critical to understand the factors that limit vaccine efficacy in advanced cancers. Here, we investigate the efficacy of neoantigen RNA-lipoplex (Neo RNA-LPX) vaccination in the preclinical tumor setting and monitor the fate of vaccine-induced T cells. We find that, in line with clinical observations, Neo RNA-LPX vaccination induces robust cytotoxic T cell responses. These T cells efficiently infiltrate tumor tissues and correlate with changes in the intratumoral myeloid compartment, characterized by an increased frequency of proinflammatory myeloid cells expressing inducible nitric oxide synthase (iNOS) and high levels of the co-stimulatory molecule CD80. This remodeling of the T cell and myeloid compartments is associated with complete regression of smaller, immature MC-38 tumors, but only delayed growth of larger, established tumors. Following cessation of therapeutic vaccination, the abundance and diversity of neoantigen-specific effector CD8 TILs rapidly declines, and the remaining CD8 T cells exhibit a phenotype resembling that of spontaneous CD8 TILs from unvaccinated mice. Single cell RNAseq analysis further revealed that this T cell contraction may be driven by a pro-apoptotic program, while surviving tumor infiltrating T cells adopted expression signatures of cellular stress and impaired activation. Importantly, recurrent weekly Neo RNA-LPX vaccination replenishes the intratumoral pool of functional neoantigen-specific CD8 T cells and improves therapeutic anti-tumor efficacy.

## Results

### Neo RNA-LPX vaccination generates long-lived neoantigen-specific CD8 T cells in blood that provide prophylactic tumor protection

To investigate the effect of neoantigen RNA-LPX (Neo RNA-LPX) vaccination in the MC-38 tumor model, we designed two mRNA decatopes, each encoding 10 unique MC-38-specific neoantigens^26,27^, which were complexed into a lipoplex formulation for intravenous injection^28^ (Fig. 1a, Supplementary Table 1). We first assessed the breadth, magnitude, quality, and longevity of neoantigen-specific CD8 T cells induced by Neo RNA-LPX vaccination in the spleen and blood of tumor-naive mice. Following three immunizations with Neo RNA-LPX (Fig. 1b), IFNg ELIspot analysis of total splenocytes revealed that 11 out of 20 neoantigens were likely immunogenic, as 10 significantly increased IFNg production and 1 demonstrated a trending increase. Secondary validation utilizing CD4-depleted or CD8-depleted splenocytes confirmed immunogenicity of 9 of the neoantigens, with 8 eliciting CD8-specific responses (M16, M2, M10, M170, M20, M143, M111, and M86) and 1 eliciting a CD4-specific response (M47) (Fig. 1c,d). As expected, a large magnitude and percentage of neoantigen-specific CD8 T cells could be detected in the blood following the second vaccination, with vaccine-induced cells comprising over 20% of the CD8 T cell compartment (Fig. 1e). On day 21 (7 days after the third vaccination), near the peak of the vaccine-induced T cell response, phenotypic analysis revealed that neoantigen-specific CD8 T cells in the blood had an activated effector phenotype. When compared to total CD8 T cells from animals that received no treatment, Neo RNA-LPX-induced neoantigen-specific CD8 T cells had decreased expression of markers associated with a stem-like or naive state (TCF1, CD62L) and increased expression of markers associated with activation, proliferation, and effector function (CD44, PD1, Ki67, and GzmB) (Fig. 1f). Importantly, neither population of CD8 T cells showed expression of TIM3 or TOX, suggesting that differentiation was not terminal and functionality remained intact (Fig. 1f). Following T cell contraction after the third immunization, a stable pool of neoantigen-specific CD8 T cells could be detected in the blood of vaccinated mice for several weeks, suggesting formation of long-lived memory cells (Fig. 1e). In support of this, we identified a pool of neoantigen-specific central memory (CD44+CD62L+) CD8 T cells in the blood that were enriched weeks after final vaccination (day 49) in comparison with the peak of response (day 19) or during contraction (day 35) (Fig. 1g). Memory formation was confirmed when boost vaccination on day 71, administered 57 days after the third vaccination (Fig. 1b), led to an expansion in the number of neoantigen-specific CD8 T cells (Fig. 1e). Importantly, vaccine-induced memory cells were found to have retained long-term functionality following MC-38 tumor challenge, as 9 out of 10 vaccinated animals were protected against tumor growth (Fig. 1h).

**Figure 1.**
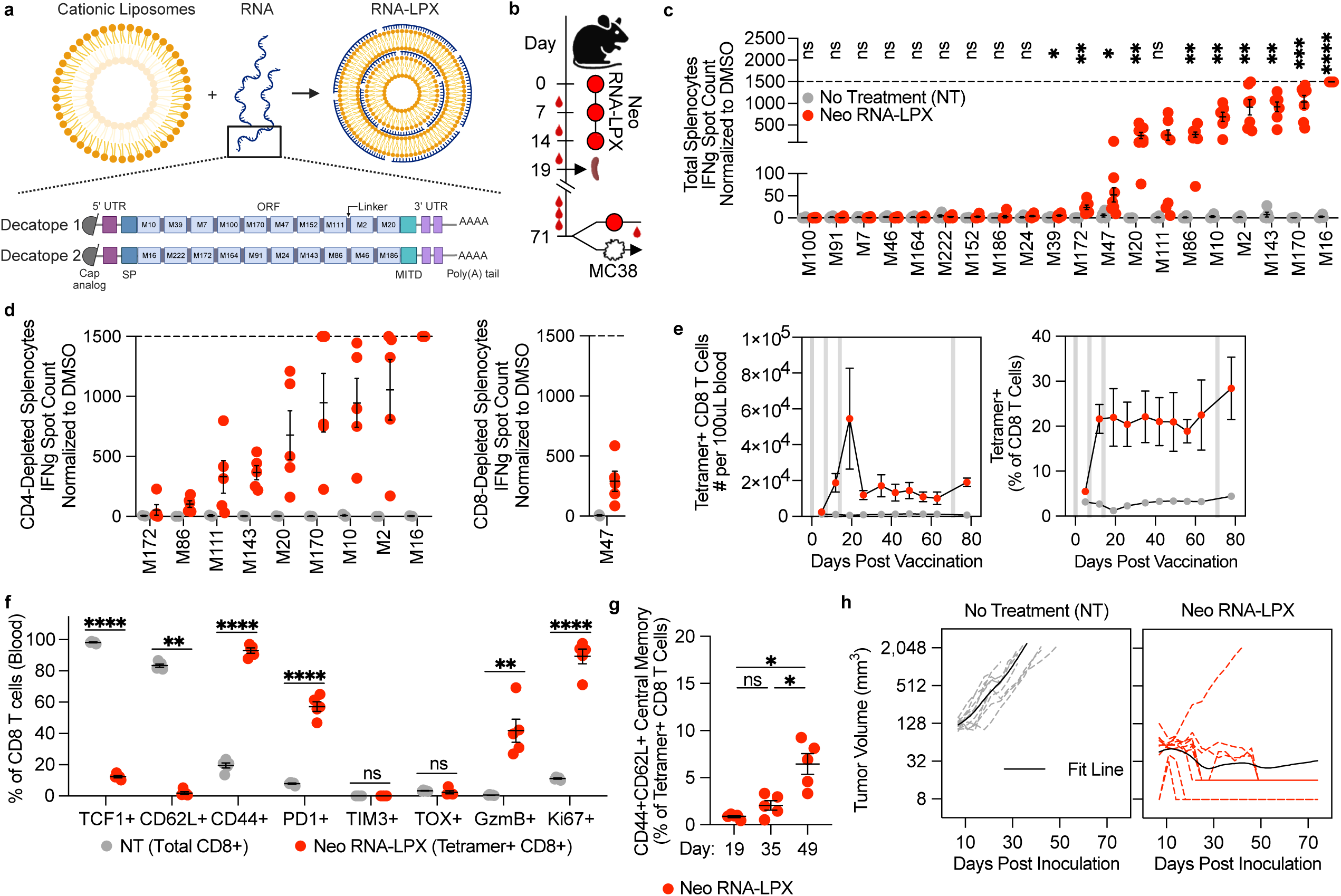
Neo RNA-LPX vaccination generates long-lived neoantigen-specific CD8 T cells in tumor-naive mice. **a**, Neo RNA-LPX vaccine composition and structure of the two decatope mRNAs, each encoding 10 different neoantigens. **b**, Mice received no treatment (NT) or were vaccinated with Neo RNA-LPX (50ug) on days 0, 7, and 14. On day 19 spleens were collected for ELIspot analysis (in **c**, **d**). T cell responses in blood were monitored following each vaccination (in **e-g**). On day 71 mice received either no treatment, an additional boost vaccination, or were challenged with 0.1e6 MC-38 tumor cells followed by monitoring of T cell responses in the blood and tumor growth, respectively (in **e**, **h**). **c**, **d**. Day 19 IFNg ELIspot analysis of total (**c**, n=5-7/group), CD4-depleted (**d**, left, n=2-5/group), or CD8-depleted (**d**, right, n=2-5/group) splenocytes following overnight stimulation with neoantigen peptides (10ug/mL) or DMSO. Dotted line depicts the upper limit of detection. **e**, Blood neoantigen-specific CD8 T cell quantification post vaccination measured by flow cytometry staining with pooled peptide:MHCI tetramers (n=4-5/group). Shaded bars indicate vaccinations. **f**, Phenotype of total CD8 T cells (NT) and neoantigen-specific CD8 T cells (Neo RNA-LPX) in the blood on day 21 after treatment initiation (n=5/group). **g**, Percentage of central memory (CD44+CD62L+) neoantigen-specific CD8 T cells in the blood of Neo RNA-LPX vaccinated mice at the indicated time points (n=5/group). **h**, Tumor outgrowth following prophylactic vaccination (n=10/group). Tumor growth curves for individual animals (dotted) and fit lines (solid) are shown. Data are presented as mean ± SEM. ns = non significant, *p<0.05, **p<0.01, ***p<0.001, ****p<0.0001.

Collectively, these results demonstrate that Neo RNA-LPX vaccination successfully generates long-lived neoantigen-specific CD8 T cells of multiple antigen specificities in the periphery that are functionally capable of recognizing tumor cells and preventing their growth in a prophylactic setting.

### Neo RNA-LPX induced T cells effectively infiltrate the tumor and exhibit an activated effector phenotype

We next evaluated the effect of Neo RNA-LPX vaccination in tumor-bearing mice. Mice with established MC-38 tumors were immunized twice with Neo-RNA-LPX followed by the collection of spleen and tumor tissues for immune cell analysis (Fig. 2a). Neoantigen-specific CD8 T cells were minimally detected in the spleen of untreated tumor-bearing mice, while they constituted approximately 25% of CD8 tumor-infiltrating lymphocytes (TILs), providing evidence that tumor-bearing mice spontaneously develop neoantigen-specific T cells that recognize tumor cells and accumulate in the tumor (Fig. 2b,c). After 2 immunizations with Neo RNA-LPX, tumor-bearing mice developed a neoantigen-specific CD8 T cell response of high magnitude in the spleen, reaching on average 38% of total CD8 T cells, along with a trending increase in total number of splenic CD8 T cells (Fig. 2b). More importantly, the total number of CD8 TILs significantly increased by 150-fold in the tumor after vaccination, with a significant increase in the frequency of neoantigen-specific CD8 T cells in comparison with untreated tumor-bearing mice (Fig 2c). Histological analysis of tumor tissues further confirmed this, demonstrating a 5-fold increase in the proportion of CD8+ TILs in Neo RNA-LPX vaccinated mice compared with untreated controls (Fig. 2d). Zonal analysis of the tumor tissues further revealed robust CD8 T cell infiltration following vaccination, with an increased abundance of CD8 TILs in both the marginal and central zones of tumors (Fig. 2e). In addition to increasing the number of CD8 TILs, Neo RNA-LPX vaccination broadened the breadth of the neoantigen-specific CD8 TILs. In untreated tumors, spontaneous T cell responses against two neoantigens, M16 and M86, were detectable, with M86 being the dominant response, as previously shown^29,30^. In contrast, Neo RNA-LPX vaccinated tumor-bearing mice showed T cell responses against 7 neoantigens, with M16 showing the dominant response (Fig. 2f), as was observed in vaccinated tumor-free mice (Fig. 1c).

**Figure 2.**
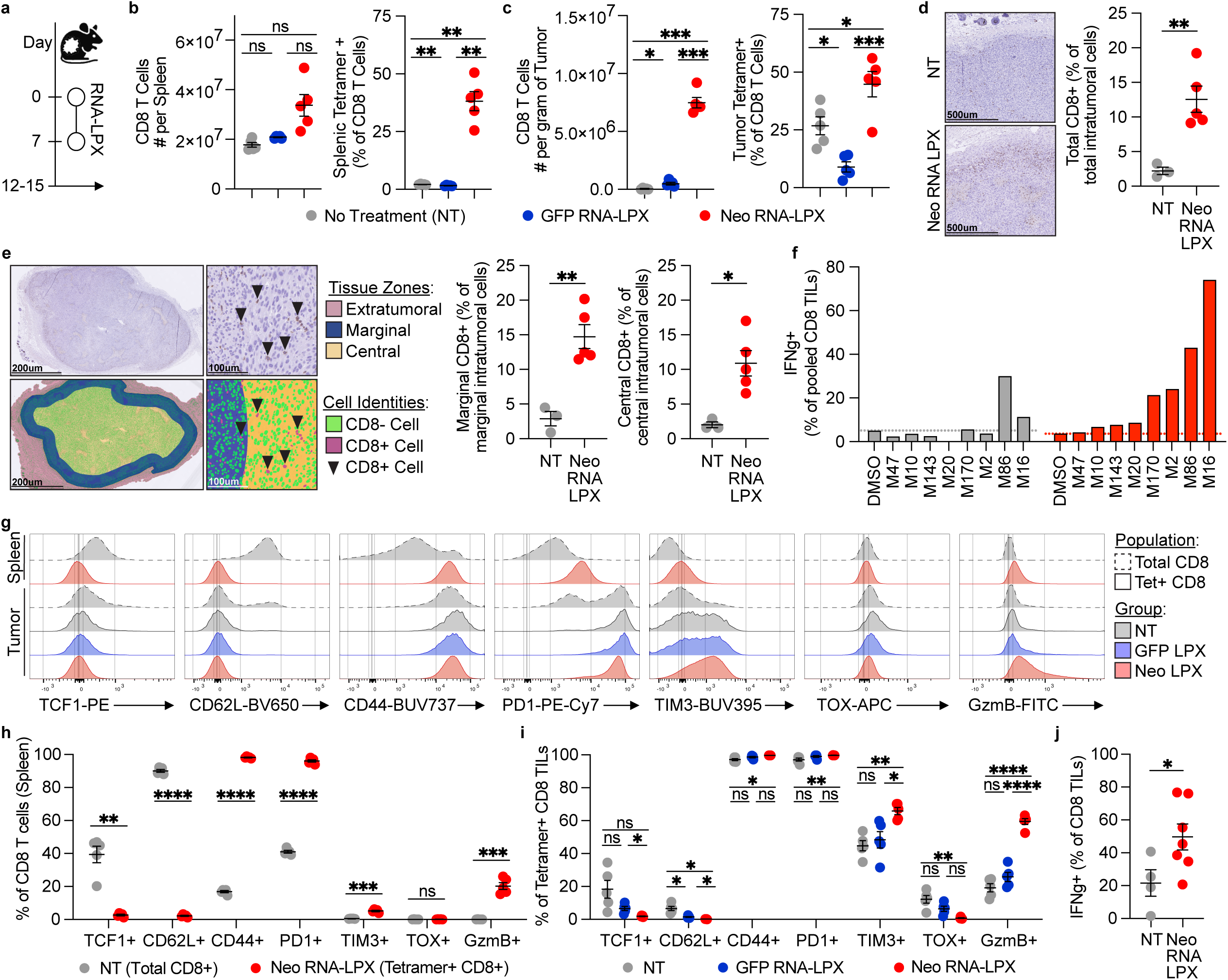
Neo RNA-LPX vaccination generates neoantigen-specific CD8 TILs with an activated and cytotoxic phenotype. **a**, Mice with established MC-38 tumors (106-258mm^3^) received no treatment (NT) or were vaccinated with Neo RNA-LPX (50ug) or control GFP RNA-LPX (50ug) on days 0 and 7. On days 12-15, tumor and spleen tissues were collected for analysis. Day 12 FACS-based quantification of total and neoantigen-specific CD8 T cells from spleen (**b**) and tumor (**c**) tissues. Day 14 histological quantification of CD8+ TILs across whole tumor (**d**) or separated based on marginal (<500um from tumor margin) and central (>500um from tumor margin) zones (**e**). **f**, Day 12 TILs were pooled from NT (n=5) and Neo RNA-LPX (n=5) vaccinated mice respectively and stimulated overnight with individual neoantigen peptides (10ug/mL) or DMSO followed by quantification of IFNg expression. **g**, Day 12 expression of T cell phenotype markers from total (dashed line) and neoantigen-specific (solid line) CD8 T cells from spleen (top) and tumor (bottom) tissues. Day 12 phenotypic composition of total and neoantigen-specific CD8 T cells in spleen (**h**) and tumor (**i**) tissues. **j**, Day 15 TILs from NT (n=4) and Neo RNA-LPX (n=7) vaccinated mice were stimulated overnight with PMA and ionomycin (1X) or DMSO followed by quantification of IFNg expression. Data are presented as mean ± SEM, representative histograms, or an individual value from pooled samples. ns = non significant, *p<0.05, **p<0.01, ***p<0.001, ****p<0.0001.

Phenotypic profiling of CD8 T cells revealed a broad array of differentiation states that varied depending on tissue localization and neoantigen specificity. In untreated tumor-bearing mice, total splenic CD8 T cells harbored high expression of TCF1 and CD62L, with only a small fraction of cells expressing CD44, indicative of a more naive state with limited antigen exposure (Fig. 2g,h). In contrast, neoantigen-specific CD8 T cells from the spleens of tumor-bearing mice vaccinated with Neo RNA-LPX expressed CD44 and low levels of PD1, TIM3, and GzmB, suggestive of an activated phenotype and recent antigen exposure, largely consistent with our findings in tumor-free mice (Fig. 2g,h, and Fig. 1f). In tumor tissues across groups, all CD8 TILs were antigen-experienced as they all expressed CD44 and PD1 with no or little expression of TCF1 or CD62L (Fig. 2g,i). While all CD8 TILs were PD1+, the level of PD1 expression was higher among neoantigen-specific CD8 TILs, indicative of prolonged TCR engagement (Fig. 2g,i). Further profiling revealed that Neo RNA-LPX vaccine-induced CD8 TILs had increased expression of TIM-3 and GzmB and reduced expression of TOX when compared with spontaneous neoantigen-specific CD8 TILs from control group mice (Fig. 2g,i). These results suggest that spontaneous neoantigen-specific CD8 TILs may be progressing toward exhaustion while neoantigen-specific CD8 TILs induced by Neo RNA-LPX vaccination appear to retain functionality. Consistent with this, ex vivo stimulation of TILs with PMA/ionomycin demonstrated enhanced expression of IFNg in CD8 TILs from Neo RNA-LPX vaccinated mice compared with the untreated control group (Fig. 2j). Vaccination with GFP RNA-LPX significantly increased the number of CD8 TILs while simultaneously decreasing the percentage of neoantigen-specific CD8 TILs (Fig. 1c), likely the result of an influx of tumor antigen irrelevant GFP-specific CD8 TILs. Notably, control GFP RNA-LPX vaccination had little to no effect on the T cell phenotype, suggesting that the RNA-LPX associated induction of systemic inflammation alone does not dramatically impact the phenotype of spontaneous neoantigen-specific CD8 TILs (Fig. 2g,i, Extended Data Fig. 1).

Collectively, these results demonstrate that Neo RNA-LPX vaccine-induced CD8 T cells are abundant in tumor tissues where they exhibit a highly activated and cytotoxic phenotype.

### Neo RNA-LPX remodels the intratumoral myeloid compartment to a more inflammatory state

Given that previous studies have shown that vaccination and immune checkpoint blockade can induce changes in intratumoral myeloid cells^31–34^, we next assessed the impact of Neo RNA-LPX vaccination on the intratumoral myeloid compartment. Neo RNA-LPX vaccination significantly enhanced the accumulation of multiple myeloid populations by approximately ∼4-6-fold, including monocytic cells (MCs), macrophages (Macs), and DCs (Fig. 3a,b). In contrast, the accumulation of neutrophils (Neus), which were present in low abundance relative to other myeloid populations, was not significantly impacted by vaccination (Fig. 3b). Phenotypic analysis further revealed that Neo RNA-LPX vaccination broadly increased the expression of iNOS across all surveyed populations, suggestive of an activated pro-inflammatory state, while expression of CD80 was largely unchanged (Fig. 3c). To better understand the functional status of iNOS+ myeloid cells in the tumor microenvironment, we further assessed CD80 expression on iNOS- and iNOS+ myeloid populations from Neo RNA-LPX vaccinated mice. Doing so revealed that iNOS+ cells also broadly had increased expression of CD80, further indicative of an activated state with enhanced capacity for co-stimulation within the tumor microenvironment (Fig. 3d). Notably, although control GFP RNA-LPX vaccination did not significantly alter the accumulation or phenotype of tumor-infiltrating myeloid populations, a trending increase in their abundance was observed (Fig. 3b). These results suggest that the systemic inflammatory response elicited by IV administration of RNA-LPX vaccines (Extended Data Fig. 1) may promote some accumulation of tumor-infiltrating myeloid cells, while myeloid cell activation may depend on the presence of activated TILs, consistent with earlier observations^28,31–34^.

**Figure 3.**
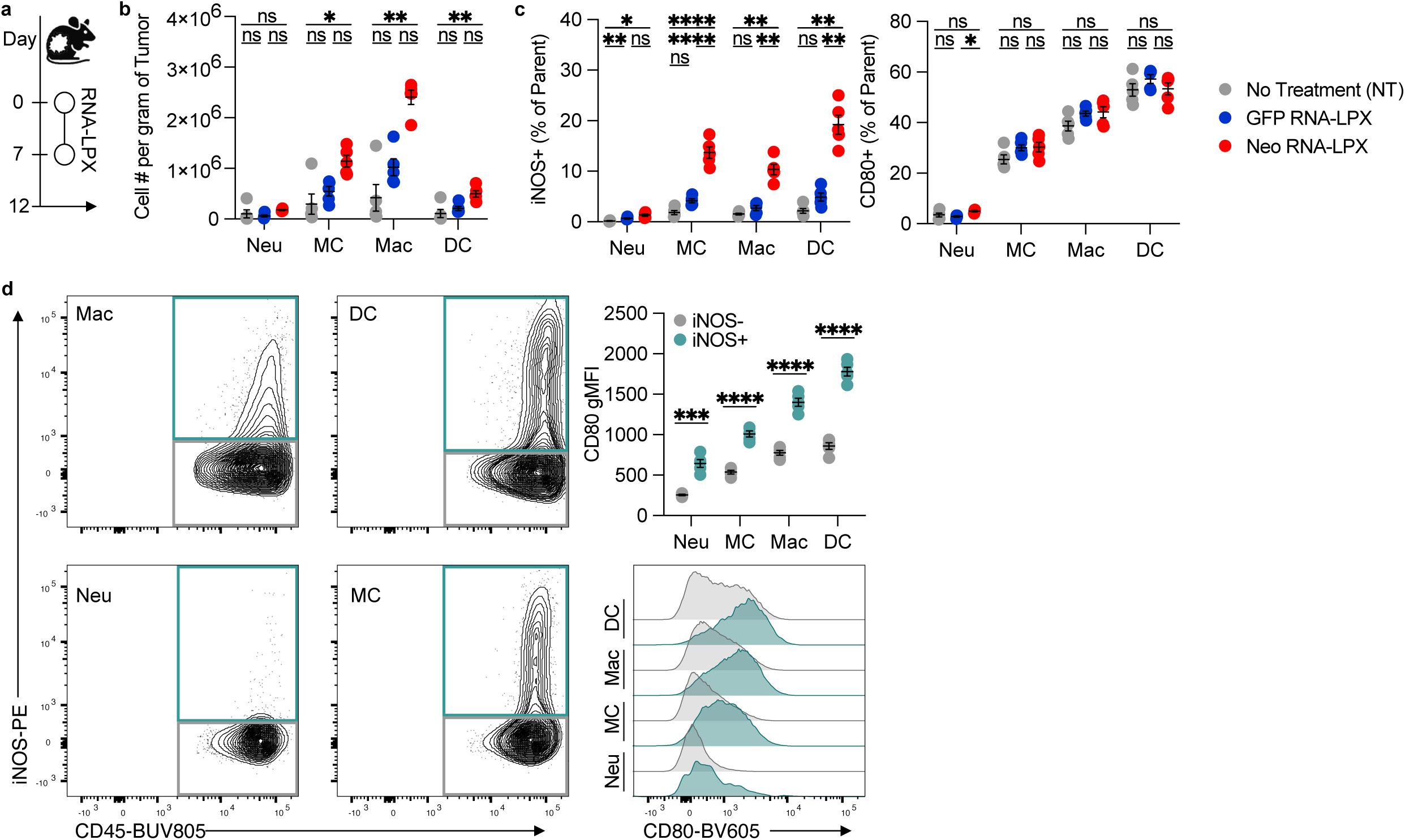
Neo RNA-LPX vaccination remodels the tumor microenvironment to a more inflammatory and stimulatory state. **a**, Mice with established MC-38 tumors (162-214mm^3^) received no treatment (NT) or were vaccinated with GFP RNA-LPX (50ug) or Neo RNA-LPX (50ug) on days 0 and 7. On day 12, tumor tissues were collected for analysis. **b**, Quantification of the number of tumor-infiltrating neutrophils (Neu; Gr1 high F480-CD11b+), monocytic cells (MC; Gr1 low F480-CD11b+), macrophages (Mac; Gr1-F480+ CD11b+), and dendritic cells (DC; Gr1-/low F480-CD11c+ MHCII high) (n=5/group). **c**, Percent of tumor-infiltrating myeloid populations expressing iNOS and CD80 (n=5/group). **d**, Neus, MCs, Macs, and DCs from Neo RNA-LPX vaccinated mice (n=5) were gated on iNOS- and iNOS+ subpopulations and expression of CD80 was measured by geometric mean fluorescence intensity (gMFI). Data are presented as mean ± SEM, representative contour plots, or representative histograms. ns = non significant, *p<0.05, **p<0.01, ***p<0.001, ****p<0.0001.

All together, our results demonstrate that Neo RNA-LPX vaccination shifts the tumor microenvironment to a more inflammatory state by not only promoting the infiltration of tumor-specific effector T cells, but also stimulating the accumulation of activated myeloid cells.

### Neo RNA-LPX vaccination mediates complete growth control of small, immature MC-38 tumors, but only delays growth of larger, established tumors

Given the capacity of Neo RNA-LPX to induce large numbers of IFNg-producing neoantigen-specific CD8 TILs and remodel the TME, we next investigated the effectiveness of vaccination at controlling tumor growth. Mice were inoculated with MC-38 tumor cells, and neoantigen vaccination was initiated on day 6, 12, or 18 after tumor inoculation to assess anti-tumor activity in tumors of different stages of development (Fig. 4a). Neo RNA-LPX vaccination initiated on day 6 or day 12 after tumor inoculation led to complete responses, with regressions seen in 15/15 and 14/15 animals respectively (Fig. 4a). Conversely, initiation of vaccination on day 18 achieved only 2/15 complete responses, with an equivalent number of spontaneous regressions seen in mice receiving no treatment, 2/25 (Fig. 4a). Importantly, while no meaningful complete regressions were seen with day 18 initiation, delayed tumor growth was evident in approximately 7/15 mice, demonstrating a clear relationship between tumor growth control and the time at which therapy was initiated (Fig. 4a). To determine whether the vaccine adjuvanticity contributed to tumor growth control, mice were treated with an irrelevant GFP RNA-LPX. Control GFP vaccination had minimal impact on tumor growth as compared with Neo RNA-LPX (Fig. 4b). This result indicates that adjuvanticity of RNA-LPX alone is not sufficient for growth control of established tumors.

**Figure 4.**
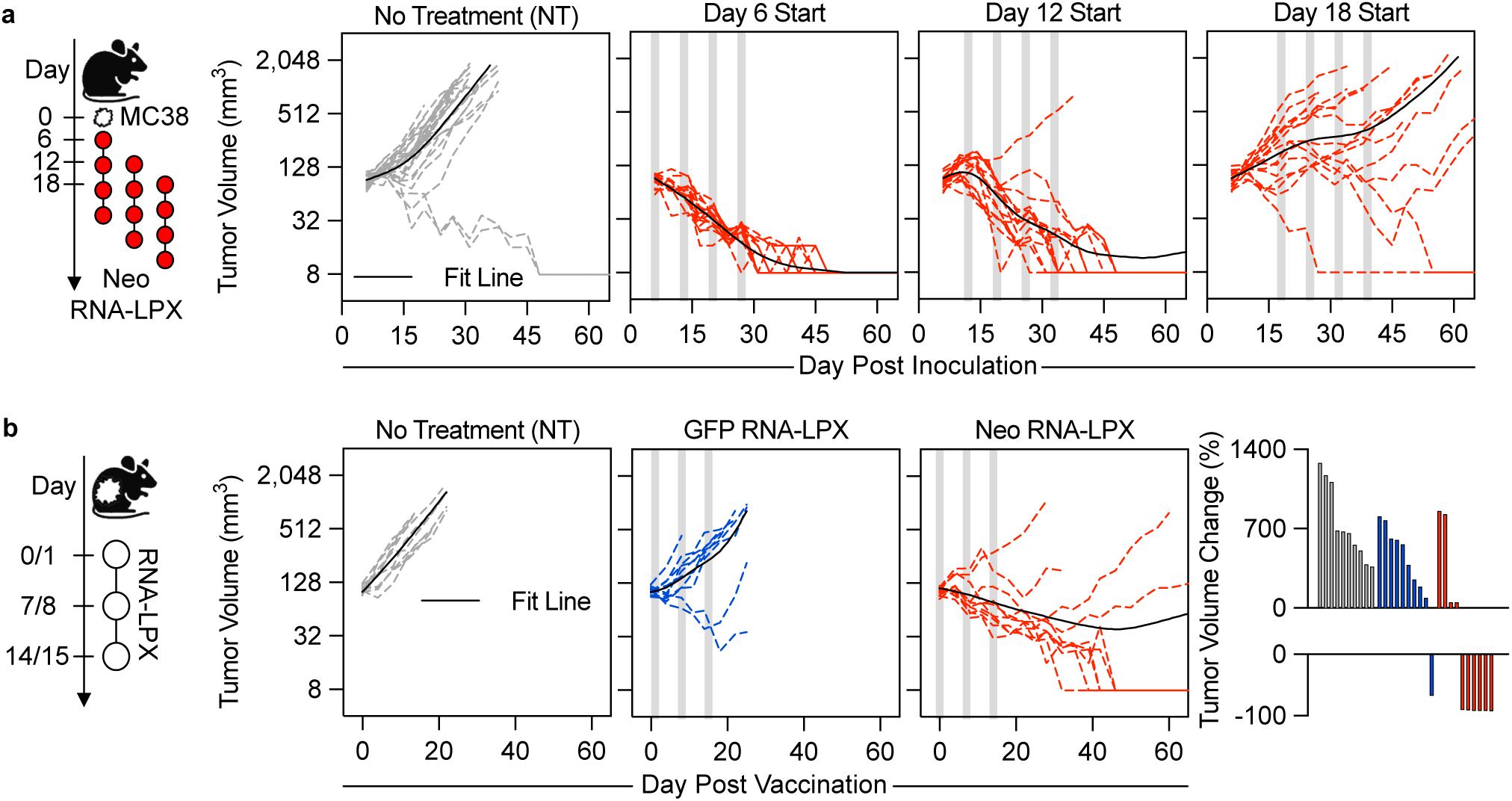
Therapeutic anti-tumor activity of Neo RNA-LPX vaccination is dependent on the stage of tumor development. **a**, Mice were inoculated with MC-38 tumor cells on day 0. Starting on either day 6 (61-105mm^3^), 12 (63-168mm^3^), or 18 (41-509mm^3^) after tumor inoculation, animals received no treatment (NT) (n=25) or began receiving the first of four total weekly doses of Neo RNA-LPX (50ug) (n=15/group). Tumor growth was monitored throughout. **b**, Mice bearing established MC-38 tumors (85-124mm^3^) received NT or were vaccinated with GFP or Neo RNA-LPX (50ug) on days 0 or 1, 7 or 8, and 14 or 15 (n=10/group). Tumor growth was monitored throughout and the percent change in tumor volume was quantified. Tumor growth curves for individual animals (dotted) and fit lines (solid) are shown.

These data demonstrate that therapeutic Neo RNA-LPX vaccination leads to the rapid elimination of immature tumors, but is insufficient to drive regression of larger, more mature tumors.

### Neo RNA-LPX-induced TILs are short-lived, with remaining TILs adapting stress responses and reverting to a pre-vaccination state

To evaluate the durability of the Neo RNA-LPX-induced CD8 T cell response in tumor-bearing mice, we performed two weekly immunizations and analyzed neoantigen-specific T cells 5 and 12 days after the final vaccination (Fig. 5a). While the number of neoantigen-specific T cells in the spleen remained stable, their number within the tumor showed a significant ∼4-fold decrease between day 5 and day 12 post-immunization (Fig. 5b). To understand the fate of the T cell response and explore any qualitative changes, we performed single-cell RNA and T cell receptor (TCR) sequencing of neoantigen-specific TILs from both vaccinated and non-vaccinated tumor-bearing mice (Fig. 5a, Extended Data Fig. 2a). In non-vaccinated tumor-bearing mice, the spontaneously developed neoantigen-specific T cell response was dominated by a T effector population (T_spontaneous), marked by expression of inhibitory receptors *Pdcd1*, *Ctla4*, *Tigit, Lag3*, but intermediate *Tox* expression, and cytolytic genes *Prf1, Gzmf*, *Gzme*, and *Gzmd* (Extended Data Fig. 2a, and Supplementary Table 2), suggesting an activation state transitioning to exhaustion, as previously observed^35^.

**Figure 5.**
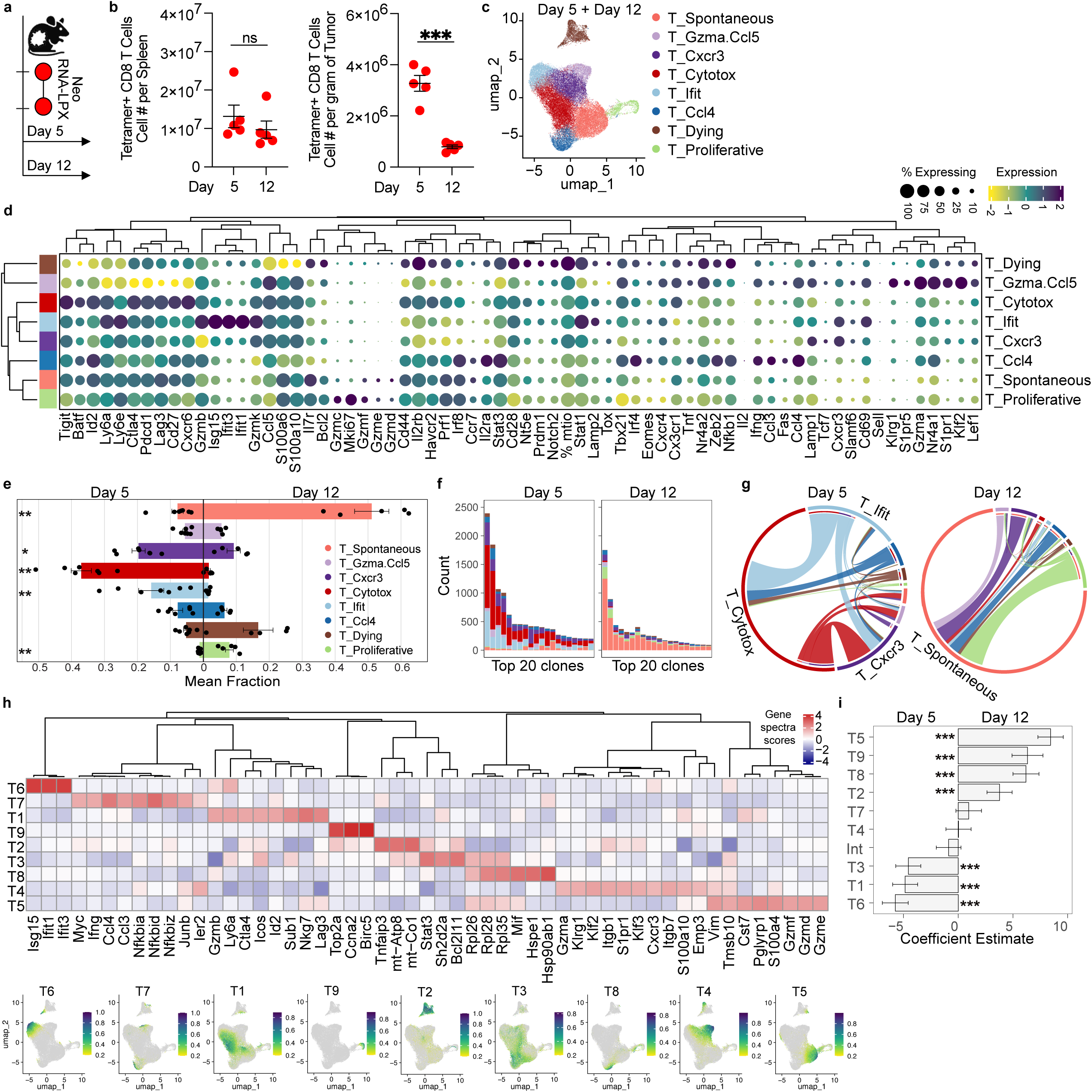
Neo RNA-LPX-induced TILs are short-lived, with remaining TILs adapting stress responses and reversion to a pre-vaccination T effector landscape. **a,** Mice with established MC-38 tumors (110-213mm^3^) received two weekly Neo RNA-LPX (50ug) vaccinations, followed by tissue collection at day 5 and day 12 post-second vaccination, followed by scRNA and scTCR sequencing of sorted neoantigen-specific CD8 TILs (n=5-7/timepoint). **b,** Quantification of the number of tetramer-positive neoantigen-specific CD8 T cells in spleen and tumor tissues at day 5 and 12 post-second vaccination. Data are presented as mean ± SEM. ns = non significant, ***p<0.001. **c,** UMAP projection of the scRNA-seq data of neoantigen-specific CD8 TILs day 5 and day 12 post-vaccination, colored by identified T cell phenotype cluster. **d,** Dotplot with z-scored expression of selected T cell phenotype markers. **e,** Barplot with the fraction of cells in each T cell phenotype cluster relative to the total number of cells per mouse. Each dot corresponds to the fraction of cells in one mouse. **f,** Stacked barplot of the top 20 expanded clones and their clonal count at day 5 and day 12 colored by the T cell phenotype. **g,** Clonal interactions among T cell phenotypes at day 5 and day 12; each link indicates shared clonotypes with line thickness denoting the interaction strength. **h,** Heatmap of z-scored gene spectra scores of selected genes from the top 15 topic genes, alongside mapping of selected topic’s usage metric onto the UMAP projection of the scRNA-seq data. **i,** Barplot of the association between cell topic modeling scores and day post-final vaccination as coefficient estimates from logistic regression analysis. Error bars represent 95% confidence intervals, and significance levels are indicated by asterisks: *p<0.05, **p<0.01,***p<0.001.

Upon Neo RNA-LPX vaccination, the neoantigen-specific T cell landscape shifted, becoming dominated by different effector populations (Fig. 5c-e, Supplementary Table 3). By day 5 post-immunization, neoantigen-specific TILs largely expanded into three dominant T cell effector phenotypes, T_Cxcr3, T_Ifit, and T_Cytotox (Fig. 5e, f). T_Cxcr3 was characterized by high expression of *Cxcr3* suggesting either recent tumor ingress or intratumoral trafficking to APCs^36^. This population exhibits intermediate expression of inhibitory receptor genes (*Pdcd1*, *Lag3*, *Tigit, Ctla4*) and *Gzmk*, but low expression *Prf1* and *Gzmb*, suggesting an early effector state poised for differentiation. T_Ifit was defined by high expression of interferon-stimulated genes (*Stat1*, *Isg15*, *Ifit3*, *Ifit1, Ly6a, Ly6e*), with elevated *Gzmb, Gzmk, Cd69,* and *Cxcr3* expression, indicating IFN-responsive, recently activated effector cells as previously described^37^. In contrast, T_cytotox is characterized by the highest expression of *Cxcr6* together with the most consolidated expression of inhibitory receptor genes (*Pdcd1*, *Tigit*, *Ctla4*, *Lag3*, *Havcr2*), suggesting repeated antigen stimulation and a highly differentiated effector phenotype. Notably, we also identified a population resembling the spontaneous T cell population found in non-vaccinated mice (Fig. 5e, Extended Data Fig. 2a-c). This population represented a minor fraction of the cells at day 5 post-immunization. Clonal network analysis revealed that on day 5, vaccine-induced effector populations (T_Cxcr3, T_Ifit, and T_cytotox) dominated the landscape with minimal overlap with the T_spontaneous population, indicating they are most likely *de novo* vaccine-primed T cells (Fig. 5g).

By day 12 post-immunization, the CD8 TIL landscape changed, with a significant reduction in the vaccine-induced effector populations, suggesting that these effector T cells may be short-lived in the tumor (Fig. 5e). Concurrently, the T_spontaneous cell population was enriched, mapping to the same clonally expanded cluster found in non-vaccinated mice (Fig. 5e, Extended Data Fig. 2c). This shift in cellular abundance was mirrored by a reduced clonal expansion (Fig. 5f). Furthermore, the majority of the clones in the T_spontaneous population did not overlap with the few remaining vaccine-induced T effector clones, indicating that these are likely not differentiated from the vaccine-induced effector clones, but rather new infiltrating clones.

To further elucidate the cellular programs driving the contraction of the day 5 T cell populations and the emergence of the day 12 T cell states, we conducted a topic modeling analysis (Fig. 5h, Extended Data Fig. 2d, Supplementary Table 4). This analysis confirmed the remodeling of the T cell landscape, with Topic 1 and Topic 6 significantly associated with day 5 T cells and largely overlapping with T_cytotox and T_Ifit, respectively, while Topic 5 and Topic 9, enriched in day 12 T cells, overlapped with T_spontaneous and T_proliferative, respectively (Fig. 5i, Extended Data Fig. 2d). We also identified cellular programs that further distinguished day 5 and day 12 TILs, reflecting transcriptional states shared across multiple effector T cell populations rather than being confined to a single population. Day 5 T cells were significantly associated with Topic 3, a cellular program characterized by the expression of Stat3 associated with cytokine responsiveness and sustained effector function^38,39^, RPL family genes indicative of high ribosomal/translational capacity, *Sh2d2a* suggestive of prolonged TCR engagement, and the pro-apoptotic gene *Bcl2l11* (Fig. 5h,i, and Supplementary Table 4). This gene expression profile indicates that a subset of day 5 T cells across phenotypes (Extended Data Fig. 2d) is characterized by a terminally differentiated effector state and prone to apoptosis. On the other hand, day 12 T cells were significantly associated with Topic 8 and Topic 2 across several T effector phenotypes. Topic 8 is characterized by the expression of *Hsp90ab1* and *Hspe1*, suggestive of stress adaptation, as well as *Mif*, which expression in tumor and myeloid cells has been linked to T cell dysfunction in tumors^40,41^. Topic 2 is characterized by *Tnfaip3,* a regulator of dysfunction in T cells^42^. These findings suggest that day 12 TILs exhibit overall a higher state of impaired activation and cellular stress. Collectively, our data demonstrate that while Neo RNA-LPX induces long-lived effector T cells that persist in the periphery, their expansion and survival is only transient in the tumor bed. Following cessation of vaccination, the T cell landscape rapidly reverts to a state resembling the pre-vaccination response in MC-38 tumors with cellular programs of T cell dysfunction and stress-adaptation.

### Recurrent vaccination replenishes proliferative and cytotoxic neoantigen-specific CD8 TILs and improves efficacy

Given the rapid decline in the abundance and transcriptional diversity of neoantigen-specific CD8 TILs following cessation of Neo RNA-LPX vaccination, we hypothesized that recurrent vaccination could replenish the population of functional TILs and strengthen anti-tumor efficacy. To test this, we first evaluated the impact of recurrent vaccination on CD8 TIL abundance and phenotype in mice that had received either 2 or 3 doses of Neo RNA-LPX (Fig. 6a). Addition of a third immunization increased the number of neoantigen-specific TILs in some mice, but due to the variability in response, the difference was not statistically significant (Fig. 6b). The fraction of neoantigen-specific TILs expressing TCF1, PD1, TIM3, or TOX, was comparable after 2 or 3 weekly immunizations (Fig. 6c). However, a third vaccination did enhance the cytotoxic and proliferative capacity of neoantigen-specific CD8 TILs as measured by increased expression of GzmB and Ki67 (Fig. 6c). Subset analysis further revealed that recurrent vaccination enhanced the fraction of both GzmB+ and GzmB-proliferating (Ki67+) neoantigen-specific CD8 TILs while the fraction of non-proliferating TILs declined (Fig. 6d). These results suggest that vaccination not only replenishes CD8 TILs through enhanced proliferation, but that a subset of these cells harbor cytotoxic potential and likely contribute to anti-tumor activity.

**Figure 6.**
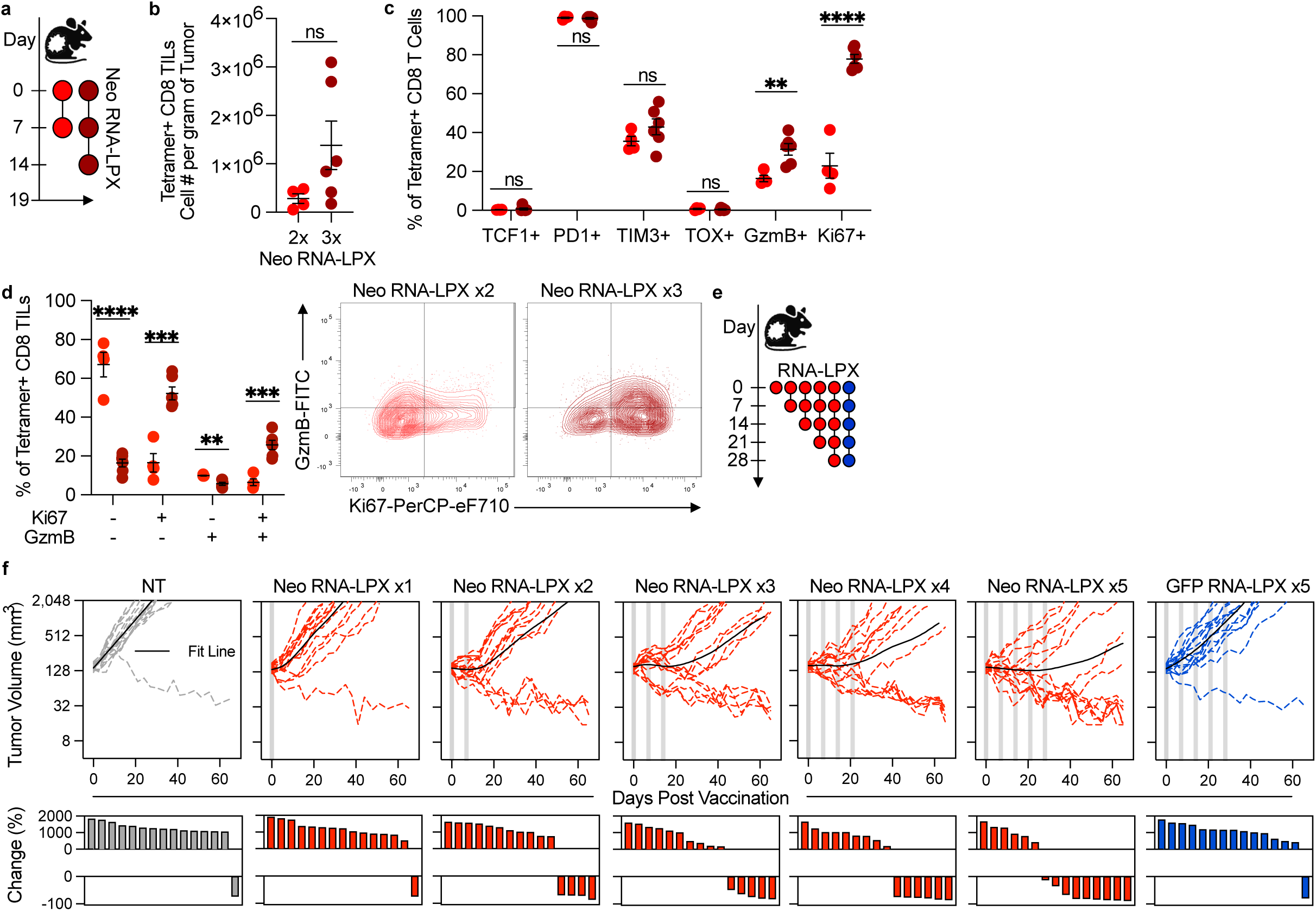
Recurrent vaccination replenishes proliferative and cytotoxic tumor-infiltrating neoantigen-specific CD8 T cells and drives anti-tumor activity. **a**, Experimental design for **b-d**. MC-38 tumor-bearing mice were vaccinated with 2 or 3 doses of Neo RNA-LPX (50ug) followed by tissue collection and analysis on day 19. Quantification of the number of neoantigen-specific CD8 TILs (**b**) and their expression of single markers (**c**) or co-expression of Ki67 and GzmB (**d**) (n=4-6/group). **e**, MC-38 tumor-bearing mice received no treatment (NT), either 1, 2, 3, 4, or 5 doses of Neo RNA-LPX (50ug), or 5 doses of GFP RNA-LPX (50ug) (n=15/group). **f**, Tumor growth was monitored throughout and the percent change in tumor volume was quantified. Tumor growth curves for individual animals (dotted) and fit lines (solid) are shown. Data are presented as mean ± SEM or representative contour plots. ns = non significant, *p<0.05, **p<0.01, ***p<0.001, ****p<0.0001.

Having established the beneficial impact of recurrent vaccination on the quality of neoantigen-specific CD8 TILs, we next assessed the impact of recurrent Neo RNA-LPX vaccination on anti-tumor efficacy. Mice with established MC-38 tumors were given up to five weekly immunizations and tumor growth was monitored (Fig. 6e). A single Neo RNA-LPX immunization had little to no impact on tumor growth, likely due to limited expansion of neoantigen-specific T cells (Fig. 6f, Fig. 1e). As the number of immunizations increased, response rates increased, with progressively increasing numbers of regressing tumors (Fig. 6f). Notably, some tumors remained stable as long as the mice were vaccinated, but progressed rapidly after the last vaccination (Fig. 6f). This effect was dependent on the neoantigen-specific T cell response, as repeated vaccination with control GFP RNA-LPX provided no benefit. Collectively, these results demonstrate that recurrent Neo RNA-LPX vaccination replenishes the pool of proliferative and cytotoxic neoantigen-specific CD8 TILs and drives increased anti-tumor activity, highlighting the importance of recurrent therapeutic RNA-LPX vaccination where established tumors are present.

## Discussion

Cancer vaccines are showing promise in early stage cancers, indicating that this approach can generate functional effector T cells capable of recognizing and killing tumor cells^1–3^. In contrast, vaccine efficacy in advanced cancers has been limited^4–7^. Thus, understanding the factors limiting this efficacy is important for developing more effective cancer vaccines and combination therapies. In this study, we investigated the efficacy of Neo RNA-LPX vaccination in the preclinical tumor setting and assessed the fate of vaccine-induced CD8 T cells. In line with clinical observations, Neo RNA-LPX vaccination generated a high quantity of long-lived polyepitopic neoantigen-specific CD8 T cells with an effector phenotype in blood^2,3,5^. Prophylactic vaccination prevented the growth of MC-38 tumors and therapeutically controlled the growth of small, immature tumors, demonstrating T cell functionality. In contrast, therapeutic Neo RNA-LPX vaccination poorly controlled the growth of large, mature MC-38 tumors despite robust tumor infiltration of vaccine-induced T cells and remodeling of the tumor myeloid compartment to a more inflammatory state. Although there was an initial accumulation of neoantigen-specific T cells in MC-38 tumors following Neo RNA-LPX vaccination, this was only transient as the abundance, diversity, and functionality of neoantigen-specific CD8 TILs rapidly declined following cessation of vaccination.

T cells induced by RNA-LPX vaccination were long-lived in the blood and spleen, indicating that their loss in large established MC-38 tumors resulted from their exposure to the TME (Fig. 5b). Five days post vaccination, single cell RNAseq analysis revealed neoantigen-specific effector CD8 T cells at different differentiation stages were found in MC-38 tumors. A dominant program (Topic 3) reflected a terminal effector state with sustained effector function through *Stat3*^38,39^ and high translational capacity (RPL genes). However, concurrent expression of the pro-apoptotic gene *Bcl2l11* (BIM) suggests a tightly controlled state in which these highly activated cells may be fated for death. In this context, external factors can tip the balance from sustained effector functions to apoptosis. The IL-21/IL-10-STAT3 axis maintains differentiation and effector function of CD8 T cells^38,43^, and the absence of these signals - whether due to lack of IL-21-producing CD4 helper cells (Tfh/Th17-like), insufficient DC or macrophage-derived IL-10, or suppression by the immunosuppressive TME - can impair effector persistence and favor apoptosis through BIM. In addition, metabolic stress can trigger apoptosis of these highly activated effector T cells as their function critically depends on sustained glycolysis, amino acid availability, and adaptation to oxidative stress^44–46^, and excessive ROS can synergize with the pro-apoptotic activity of BIM (*Bcl2l11*) to drive apoptosis^47^. Our data shows that the few neoantigen-specific CD8 T cells remaining at day 12 were enriched for markers of dysfunction and metabolic stress as shown by expression of *Tnfaip3* (Topic 2), *Mif*, *Hsp90ab1*, and *Hspe1* (Topic 8) (Fig. 5h), and suggests that metabolic stress could at least partially contribute to T cell functional impairment and trigger apoptosis of highly activated effector TILs. Consistent with this, 12 days post vaccination, the T cell landscape appears to reverse to its original state, with the phenotypes of the d12 T cells largely mirroring the ones in unvaccinated mice, including a majority of T cells belonging to the spontaneous unvaccinated TIL population (T_spontaneous). Furthermore, the majority of the T_spontaneous T cell clones do not overlap with those of the vaccine-induced phenotypes. These results suggest that d12 T cells are not directly vaccine-induced, rather they may have been spontaneously primed before vaccination or indirectly primed after vaccination through epitope spreading. These TILs express high Il7r and Ccl5 suggestive of some effector activity and persistence. However, they express reduced granzyme B compared to vaccine induced T cells (Fig. 2g) and are insufficient for tumor growth control. Given that granzyme B production is dependent on glucose utilization, this impaired effector function may be triggered to favor persistence in the nutrient poor tumor milieu^48^. Our results further suggest that the fate of vaccine-induced T cells differs from spontaneous T cells. What differentiates these spontaneous T cells from the vaccine induced T cells and why they appear to have a different fate in the tumor will require additional studies.

Neo RNA-LPX vaccination remodeled the intratumoral myeloid compartment, not only increasing the number of monocytic cells, macrophages, and DCs, but also enhancing their proinflammatory functions, as evidenced by the higher fraction of iNOS expressing cells with elevated expression of the costimulatory molecule CD80. Remodeling of the myeloid compartment was not the result of vaccine adjuvanticity caused by systemic proinflammatory cytokine release following intravenous administration of RNA-LPX^28,49^, as the control GFP RNA-LPX vaccine had little impact on the intratumoral myeloid compartment (Fig. 2b,c). Rather, it is the active intratumoral neoantigen-specific T cells that seem to be responsible for the myeloid remodeling, most likely through the combined secretion of effector molecules, like IFNg and TNFa, and subsequent generation of damage-associated molecular patterns (DAMPs) released for dying tumor cells^31–34,50^. Our results contrast with Baharom et al. who previously showed that intravenous administration of a self-assembling nanoparticle vaccine (SNAPvax), composed of neoantigen peptides linked to a toll-like receptor (TLR) 7/8 agonist, induced remodeling of the TME in a type I IFN-dependent manner^32,51^. However, a major notable difference is that SNAPvax distributes to the tumor where it is taken up by myeloid cells, while RNA-LPX, which is composed of larger particles (200-400nm), has poor intratumoral penetration (data not shown). While our results suggest favorable remodeling of the myeloid compartment by Neo RNA-LPX, this shift was insufficient to maintain vaccine-induced T cell function and survival in tumors.

Previous work has shown that intratumoral DCs produce interleukin 12 (IL-12) upon sensing of IFNg produced by T cells. In turn, IL-12 enhances CD8 T cell activation and function. This feedback loop allows for amplification of the immune response and is essential for effective anti-tumor immunity after anti-PD-1 treatment^52^. However, IFNg and IL-12 alone may not be sufficient for maintaining T cell function in the tumor. Antigen cross-presentation by tumor-resident DCs to T cells is also likely required. Indeed, we recently reported that RNA-LPX-induced T cells, both tumor antigen matched and unmatched controls, traffic efficiently to the tumor after adoptive transfer, but only tumor antigen matched T cells exhibited sustained effector function through clustering with tumor-resident cDC1s^53^. Vaccination broadens the breadth of the tumor neoantigen-specific T cell response by delivering tumor antigens to DCs outside the tumor, but many vaccine-targeted neoantigens may not be effectively cross-presented by DCs in tumors, preventing sustained T cell support in the TME. Indeed, we observed spontaneous T cell responses to only two neoantigens in unvaccinated mice (Fig. 2f). Additionally, CD4 T cells provide critical help to CD8 T cells in the TME through the secretion of cytokines such as IL-21 which promotes CD8 T cells survival and function^38^, and the formation of CD4 T cell/DC/CD8 T cell triads appears to be associated with superior CD8 T cell responses and anti-tumor efficacy^14,15,54,55^. Although Neo RNA-LPX induced a CD4 T cell response, it was limited to one neoantigen and may not be sufficient to support the large vaccine-induced CD8 T cell response. Alternatively, the number or the maturation status of the intratumoral DCs may be inadequate. Additional studies will be needed to determine why RNA-LPX vaccine-induced functional effector T cells are not maintained in the TME.

Finally and most importantly, vaccine-induced T cell loss in the tumor bed could be mitigated by repeated vaccination, which sustained a pool of activated vaccine-induced T cells in the periphery, enabling the replenishment of functional T cells in the tumor and improved antitumor efficacy (Fig. 6). Our results suggest that continuous immunization may be a critical factor for vaccine efficacy in patients and that combination strategies favoring improved T cell fitness and survival in the tumor microenvironment will improve vaccine activity in the metastatic setting.

## Methods

### Mice

Animals were maintained in accordance with the Guide for the Care and Use of Laboratory Animals (National Research Council 2011). Genentech is an AAALAC-accredited facility and all animal activities in this research study were conducted under protocols approved by the Genentech Institutional Animal Care and Use Committee (IACUC). Wildtype C57BL/6NCrl female mice were purchased from Charles River and used for experiments when they were 6-10 weeks of age. Mice were maintained in a specific-pathogen-free facility, in individually ventilated cages within rooms maintained on a 14:10-hour, light:dark cycle. Animal rooms were temperature and humidity-controlled, at 68–79 °F (20.0 to 26.1 °C) and 30–70% respectively, with 10 to 15 room air exchanges per hour.

### RNA-LPX vaccine formulation and administration

Unmodified RNAs encoding MC-38 neoantigens were generated at Genentech. Liposomes were composed of a 2:1 molar ratio of the cationic lipid DOTMA and the helper lipid DOPE. Liposomes were complexed with negatively charged RNA at a charge ratio (+):(−) of 1.3:2 in saline to yield negatively charged RNA-LPX. RNA-LPX (50ug) vaccines were administered intravenously every 7 days, with the total number of vaccinations being indicated in each figure.

### MC-38 tumor model use and analysis

The MC-38 tumor cell line was a kind gift from R. Offringa, Leiden University Medical Center, Netherlands. Upon receipt, the cell line was confirmed to be free of mycoplasma contamination and low passage stocks were created for long-term use. After culturing and expansion, early passage cells were used for subsequent animal studies. Cells were maintained in RPMI1640 supplemented with 10% FBS, and 1% L-glutamine and passaged every 3-4 days. For in vivo tumor inoculation, cells were resuspended in a 1:1 mixture of HBSS and Matrigel and injected into the right hind flank at a concentration of 0.1e6 cells per 100uL per animal. Following inoculation, tumor growth was monitored via digital caliper measurements and tumor volumes were calculated using perpendicular length and width measurements and the following formula: volume = 0.5 x length x width^2^. Analyses and comparisons of tumor growth were performed using a package of customized functions in R (Version 4.1.0: Fig. 1h and Fig.4b, Version 4.3.3: Fig. 6f, and Version 4.4.2: Fig. 4a, The R Foundation), which integrates software from open source packages (e.g., lme4, mgcv, gamm4, multcomp, settings, and plyr) and several packages from tidyverse (e.g., magrittr, dplyr, tidyr, and ggplot2)^56^. Briefly, as tumors generally exhibit exponential growth, tumor volumes were subjected to natural log transformation before analysis. All raw tumor volume measurements less than 8mm^3^ were judged to reflect complete tumor absence and were converted to 8mm^3^ prior to natural log transformation. Additionally, all raw tumor volume measurements less than 16mm^3^ were considered miniscule tumors too small to be measured accurately and were converted to 16mm^3^ prior to natural log transformation. The same generalized additive mixed model was then applied to fit the temporal profile of the log-transformed tumor volumes in all study groups with regression splines and automatically generated spline bases. This approach addresses both repeated measurements from the same study subjects and moderate dropouts before the end of the study.

### Tissue collection and processing

At the indicated time points blood, spleen, and/or tumor tissues were collected from mice for use in downstream assays. Whole blood was collected in heparin-coated tubes from the retroorbital sinus. Red blood cells were lysed with ACK lysis buffer, resuspended in FACS buffer, and the resulting cell suspensions were used for downstream cell-based assays. Spleens were collected in PBS and then physically and enzymatically digested at room temperature by crushing the spleens through a 70um filter into Cell Dissociation Buffer (Gibco) containing 12.5ug/mL Liberase TM (Sigma, Roche) and 0.1mg/mL DNase I (Sigma, Roche). Following 15 minutes of digestion, red blood cells were lysed with ACK lysis buffer, cells resuspended in FACS buffer or RPMI-1640 complete media, and the resulting cell suspensions were used for downstream cell-based assays. Tumors were collected in PBS, weighed, and then physically and enzymatically digested by mincing with scissors followed by digestion for 20 minutes at 37°C with agitation (Miltenyi gentleMACS Octo Dissociator with Heaters). Enzymatic digestion media was composed of Dulbecco’s Modified Eagle Medium (DMEM) with high glucose, 5% fetal bovine serum (FBS), 10mM HEPES, 2mg/mL Collagenase D (Sigma, Roche), and 0.1mg/mL DNase I (Sigma, Roche). Following digestion, cells were filtered (100um), resuspended in 44% percoll, centrifuged to enrich for leukocytes, resuspended in FACS buffer or RPMI-1640 complete media, and the resulting cell suspensions were used for downstream cell-based assays.

### ELIspot

Following the processing of spleen tissues into single cell suspensions, ELIspot assays were performed. Magnetic microbead-based depletion of splenocytes was carried out using the EasySep Mouse CD4 and CD8a Positive Selection II Kits (STEMCELL Technologies) with the RoboSep-16 automated cell separator (STEMCELL Technologies) according to the manufacturer’s protocol in order to obtain CD4-depleted and CD8a-depleted splenocytes. The mouse IFNg ELIspot kit (R&D Systems) was used according to the manufacturer’s protocol in conjunction with 0.5e6 total, CD4-depleted, or CD8-depleted splenocytes stimulated overnight with 10ug/mL of individual neoantigen peptides (Supplemental Table 1) or DMSO. Spot counts were obtained with an automatic ELIspot reader instrument (AID). Spot counts were normalized to DMSO control via background subtraction.

### Ex vivo functional assays

Following the processing of tumor tissues into single cell suspensions, cells were individually stimulated with 1X PMA/ionomycin (eBioscience Cell Stimulation Cocktail 500X, ThermoFisher Scientific) or cells from similarly treated mice were pooled and stimulated with individual neoantigen peptides (10ug/mL) (Supplementary Table 1). After 1 hour of stimulation, a mixture of human IL-2 (50U/mL, MilliporeSigma) and the GolgiPlug (1uL/mL, BD Biosciences) and GolgiStop (0.66uL/mL, BD Biosciences) Protein Transport Inhibitors were added and stimulation continued overnight. Cells were then collected and CD8 T cell expression of IFNg was measured via flow cytometry.

### Flow Cytometry

Following the processing of tissues into single cell suspensions, FACS analysis was performed. Cells were first stained with a fixable viability dye and Fc receptors blocked in PBS followed by staining with peptide:MHC tetramers in FACS buffer. Biotinylated peptide:MHC monomers were made at Genentech as previously described and then manually tetramerized using streptavidin-fluorophore conjugates^57^. Cells were then stained with fluorophore-conjugated antibodies for surface targets. For detection of intracellular targets, cells were fixed and permeabilized using the eBioscience Foxp3/Transcription Factor Staining Buffer Set (ThermoFisher Scientific) or the Cytofix/Cytoperm Fixation/Permeabilization Kit (BD Biosciences) according to the manufacturers protocol followed by staining with fluorophore-conjugated antibodies. Fluorescence of stained cells was measured using a BD FACSymphony flow cytometer and the resulting data was analyzed using FlowJo (Version 10). A detailed list of FACS reagents can be found in Supplementary Table 5.

### Histopathology and image analysis

Formalin-fixed, paraffin-embedded sections were deparaffinized in xylene and then rehydrated through a graded series of alcohols, ending with deionized water. Sections underwent antigen retrieval using DAKO Target Retrieval pH6 (#S1699, DAKO, Agilent Technologies) for 20 min in a water bath at 99°C and then were cooled for 20 minutes. Endogenous peroxidase activity was quenched for 4 minutes with 3% H_2_O_2_ diluted in 1X PBS. Immunohistochemistry was performed on a Thermo Scientific Autostainer 360, with sections incubated in Rabbit anti-CD8a antibody (clone 21E3.TRANS, Genentech) diluted to 1 ug/mL in 3% BSA in 1X PBS for 60 minutes at room temperature. Rabbit Monoclonal IgG (clone DA1E, Cell Signaling Technologies, Cat #3900S) diluted to 1 ug/mL in 3% BSA in 1X PBS served as an isotype control. Sections were then incubated in Powervision polymer-HRP anti-Rb (Leica, Cat PV6119) for 30 minutes, followed by DAB (Thermo Scientific, Cat #1855910) for 5 minutes. Tissues were counterstained with Mayer’s Hematoxylin (Cat.# L-756-1A, Rowley Biochemical Inc., Danvers, MA) and Bluing reagent (Cat. #7301, Thermo Scientific, Cheshire, WA) for 4 minutes each. Sections were then dehydrated through a graded series of alcohols, transferred to xylene, and covered with glass coverslips.

CD8a IHC stained slides were scanned on a Hamamatsu NanoZoomer S360 whole slide imager (C13220-01) at a final magnification of 20x. The tumor margin was digitally annotated by a pathologist. The resulting scanned images were converted to OME-Zarr^58^ and subsequently analyzed using Python (v3.8 and 3.9) along with various analysis packages, including Dask and others^59,60^. SLURM^61^ was utilized as an efficient workload and job scheduler. For the primary image analysis workflow, computational resources allocated per slide included 4 CPUs, 64 GB of RAM, and a NVIDIA Quardo P6000 GPU. Cell segmentation was performed using a grayscale Cellpose 2.0 cyto2 model^62^ retrained via human-in-the-loop on nine 224×224 image patches derived from a previously curated dataset containing CD8a/DAB-positive cells^63^. DAB positivity thresholds were determined using a combination of HSV (hue, saturation, value) and blue-channel image normalization techniques^64^. A cell was classified DAB-positive if more than 50% of its segmented area contained DAB signal. Area segmentations were calculated through a combination of pathologist-defined regions of interests (ROIs) and Otsu thresholding. Ribbon areas were identified by applying a distance transform and thresholding based on a predefined distance.

### Statistical Analysis

Shapiro-Wilk test was used to assess normality followed by use of appropriate parametric or non-parametric statistical tests. Statistical significance between two groups was determined using parametric unpaired t-test or nonparametric Mann-Whitney test. In cases of unequal variances, as determined by the F test, unpaired t-test with Welch’s correction was used. Statistical significance between more than two groups was determined using parametric one-way ANOVA with Tukey’s multiple comparisons test or non-parametric Kruskal-Wallis test with Dunn’s multiple comparisons. In cases of unequal variances, as determined by the Brown-Forsythe Test, Welch ANOVA with Dunnett T3 multiple comparisons test was used. The results of statistical tests were indicated as follows: ns = non significant, *p<0.05, **p<0.01, ***p<0.001, ****p<0.0001.

### Single-cell RNA Sequencing

Tumor tissues were processed with the Mouse Tumor Dissociation Kit (Miltenyi) using the gentleMACS Octo Dissociator with Heaters (Miltenyi) and 37C_m_TDK_1 program according to the manufacturer’s protocol. Following digestion, cells were filtered (70um), resuspended in 44% percoll, centrifuged to enrich for leukocytes, resuspended in FACS buffer, and the resulting cell suspensions were FACS stained as described above and neoantigen-specific CD8 TILs were sorted for analysis. Briefly, cell suspensions were stained with a viability dye and Fc receptors blocked followed by staining with oligonucleotide-barcoded peptide:MHC tetramers for the M2, M10, M16, M86, M111, M143, M170, and M172 neoantigens. Biotinylated peptide:MHC monomers were tetramerized using streptavidin conjugated with the PE fluorophore and TotalSeq-C oligomer (Biolegend). Cells were then stained with fluorophore-conjugated antibodies for surface targets and neoantigen-specific (tetramer+) CD8 TILs were sorted (Live CD45+ CD19-CD90.2+ CD4-CD8+ tetramer+) using the BD FACSAria cell sorter. A detailed list of FACS reagents can be found in Supplementary Table 5. Single cells were partitioned into Gel Bead-in-Emulsions (GEMs) using the Chromium single cell 5’ chip K on a Chromium controller (10X Genomics), followed by cell lysis and barcoded reverse transcription of mRNA. Paired single cell 5’ gene expression and VDJ sequence libraries were generated according to manufacturer’s instructions using the following kits: Chromium Next GEM Single Cell 5’ Kit v2 (1000263), Chromium Single Cell Mouse TCR Amplification Kit (1000254), Library Construction Kit (1000190), 5’ Feature Barcode Kit (1000256), Dual Index Kit TN Set A (1000250) and Dual Index Kit TT Set A (1000215) (10X Genomics). Gene expression libraries were sequenced on an Illumina NovaSeq 6000 instrument at read lengths of 28×90+10+10 and a sequencing depth of 20k per cell. scTCRseq libraries were sequenced on an Illumina NovaSeq 6000. FASTQ files of gene expression and TCR libraries were generated and analyzed with CellRanger (v7.1.0) using the GRCm38 mouse reference genome and VDJ GRCm38 5.0.0 for alignment, respectively.

### scRNA-seq processing and analysis

The CellRanger output filtered gene count matrices were integrated into one seurat object and filtered for cells with more than 300 and less than 6000 features, less than 5% mitochondrial and 25% globin transcripts. Feature counts were normalized per cell, scaled by 10,000, and log-transformed. T-cell receptor genes were excluded from further analysis. The top 2000 highly variable features were identified using variance stabilizing transformation, followed by data scaling, principal component analysis, and Uniform Manifold Approximation and Projection (UMAP) dimensionality reduction with 30 principal components. Clustering was performed using shared nearest neighbors and graph-based methods. Clusters suspected of cell contaminants with expression of *Cd74*, *Xist*, *Hba-a1*, *Hba-a2* were removed and data reprocessed as described above. T cell phenotypes were characterized at a clustering resolution of 0.5 using a combination of unbiased cluster marker analysis using Seurat FindAllMarkers and supervised analysis of known and published T cell phenotype and function gene expression. Analysis of T cell phenotype fractions was conducted by calculating the proportion of cells relative to the total cell count in each mouse. The MC-38 non-vaccinated single-cell RNA-seq datasets from this study (n=2) were integrated with the isotype control mice of a previously published study^65^ (n = 3), both containing CD8 tetramer-positive cell populations. Integration was performed using Harmony (v1.2.0, variable features = 2000, theta = 1). Clustering was performed as described above and T cell phenotypes were annotated at a resolution of 0.6. To compare the MC-38 untreated CD8+ T cell phenotypes with the MC-38 Neo RNA-LPX-induced CD8+ T cell phenotypes, cluster expression centroids of the Neo RNA-LPX-induced clusters were calculated using the trimmed mean of the normalized expression values and then log2 TPM transformed. The centroids were subsequently used for label transfer of the MC-38 untreated data using SingleR, based on the similarity of the expression to the centroids, followed by normalization of the resulting contingency tables to obtain labeled cluster proportions. Topic modeling was carried out using the cNMF python package (v1.5.4), with number of topics ranging from 5 to 13. The optimal number of topics was selected based on consensus stability, modeling error, and effective representation of cell clusters. For downstream analysis, gene spectra scores and the topic usage matrix from cNMF were utilized; logistic regression was applied to the topic usage matrix to assess associations between topics and days post-last vaccination, with coefficient estimates and significance levels extracted for visualization. Statistical significance of differences in T cell phenotype fractions and topic usage within T cell phenotypes across days post–last vaccination was assessed using the Wilcoxon rank-sum test. Significance levels were defined as follows: *P ≤ 0.05, **P ≤ 0.01, ***P ≤ 0.001, and ****P ≤ 0.0001.

### scTCR-seq processing and analysis

The CellRanger filtered contig annotation matrices were processed using scRepertoire v2.0.4 by combining the TCR sequences, stripping the barcodes and adding the TCR information to the merged seurat object processed as described above. Clonal assignments were made using the clone nucleotidesequence (CTnt). For clonotype analysis, cells without an assigned clonotype, cells with more than one TRB and more than two TRA were removed. To compute clonal interactions and capture shared phenotypes across T cell clones, a phenotype pair matrix was computed by summing x_k_*x ^T^ over all clones, where x_k_ represents the vector of phenotype counts for clone k, and x_k_*x ^T^ represents the outer product of the vector x_k_ with its transpose x_k_ ^T^, effectively measuring the shared presence of clonotypes between pairs of phenotype clusters. The final phenotype pair matrix was visualized as ChordDiagrams using the circlize package v0.4.16.

## Data Availability

Single-cell RNA and TCR-sequencing data generated in this study will be deposited in the Gene Expression Omnibus database and accession numbers will be made available upon manuscript acceptance.

## Code Availability

No new algorithms were developed as part of this study.

## Acknowledgements

We would like to thank Thomas Wu for his invaluable feedback and guidance on the single cell sequencing data analysis. We also thank Thomas Wu, Mathias Vormehr, and Soyoung Oh for their comments on the manuscript. We would like to thank the following groups at Genentech for their research related support: Lab Animal Resources (LAR), the Institutional Animal Care and Use Committee (IACUC), members of the Department of Protein Chemistry, especially Martine Darwish and Maria Lorenzo, the Research Flow Cytometry Core Lab, the In Vitro Cell Culture (IVCC) Lab, and the Research Pathology Core Labs, especially Allison Huynh, Jessica Mills, and Erick Pata.

## Author Contributions

Conceptualization, Writing - original draft, review, and editing: JTG, MH, JMS and LD. Formal analysis and Visualization: JTG and MH, DD, KM, and AAL. Investigation: JTG, SL, VJ, LL, MJM, TNJ, YO, DD, KM, AAL, and AT. Software; MH, KM, and JHB. Methodology and Resources: DD, KM, AAL, AMN, ECF, and AC. Project administration: CCC. Supervision: IM, US, JMS, and LD.

---

**Supplemental Figure 1.**
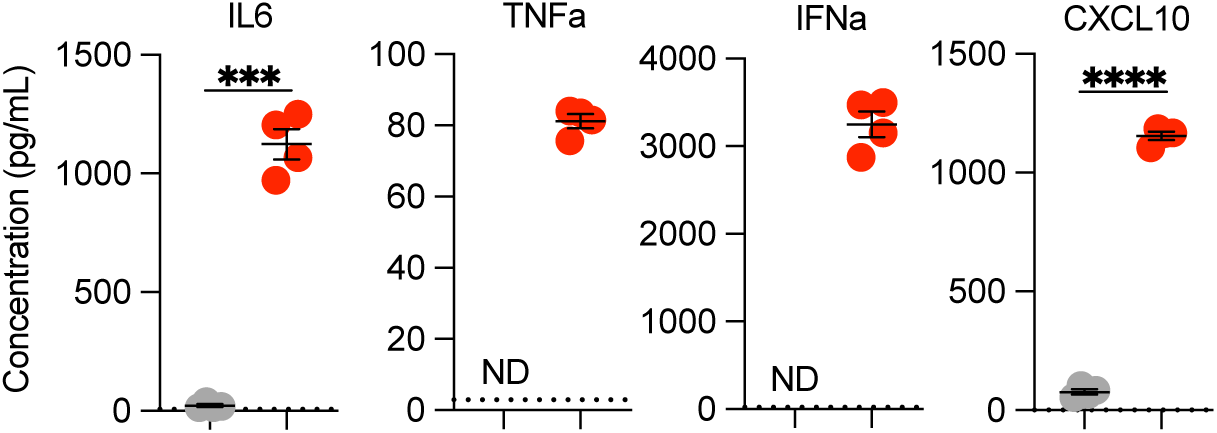
Neo RNA-LPX vaccination drives a systemic inflammatory cytokine response. Mice received no treatment (NT) or were vaccinated with Neo RNA-LPX (50ug) and 6 hours later blood was collected from the retroorbital sinus and serum was isolated. Serum cytokine and chemokine concentrations were quantified using the bead-based mouse ProcartaPlex immunoassay (ThermoFisher). Data are presented as mean ± SEM. Dotted lines represent the lower limit of detection for each cytokine. ND = not detected, ***p<0.001, ****p<0.0001.

**Supplemental Figure 2.**
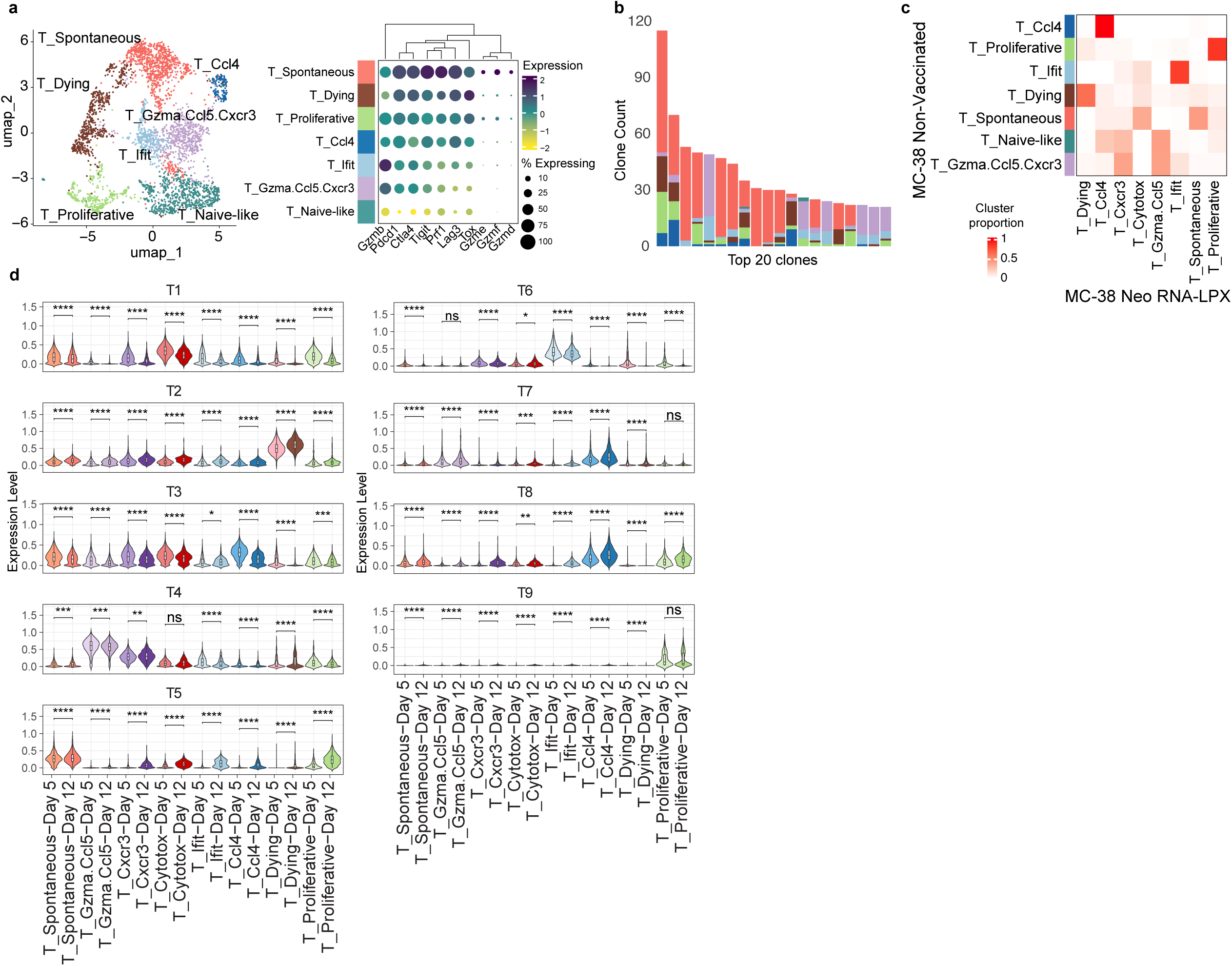
**a,** UMAP projection of scRNA-seq data of neoantigen-specific tumor CD8+ T cells of non-vaccinated MC-38 tumors, colored by identified T cell phenotype cluster. **b,** Stacked barplot of the top 20 expanded clones colored by the T cell phenotype. **c,** Heatmap showing the proportions of MC-38 non-vaccinated cluster labels mapped to the MC-38 Neo-LPX centroids. **d,** Stacked violin plots with the topic usage scores across T cell phenotypes at Day 5 and Day 12 post-last vaccination. ns = non significant, *p<0.05, **p<0.01, ***p<0.001, ****p<0.0001.

